# Precise Estimation of In Vivo Protein Turnover Rates

**DOI:** 10.1101/2020.11.10.377440

**Authors:** Jonathon J. O’Brien, Vikram Narayan, Yao Wong, Phillip Seitzer, Celeste M. Sandoval, Nicole Haste, Megan Smith, Ramin Rad, Aleksandr Gaun, Adam Baker, Matthew Kukurugya, Baby Martin-McNulty, Chunlian Zhang, Ganesh Kolumam, Carmela Sidrauski, Vladimir Jojic, Fiona McAllister, Bryson Bennett, Rochelle Buffenstein

**Affiliations:** Calico Life Sciences LLC; Department of Molecular and Cell Biology, University of California Berkeley; Calico Life Sciences, LLC

**Author notes:** Correspondence to Jonathon O’Brien, Bryson Bennett or Rochelle Buffenstein. These authors jointly supervised this work (Bryson Bennett, Rochelle Buffenstein).

## Abstract

Isotopic labeling with deuterium oxide (D_2_O) is a common technique for estimating *in vivo* protein turnover, but its use has been limited by two long-standing problems: (1) identifying non-monoisotopic peptides; and (2) estimating protein turnover rates in the presence of dynamic amino acid enrichment. In this paper, we present a novel experimental and analytical framework for solving these two problems. Peptides with high probabilities of labeling in many amino acids present fragmentation spectra that frequently do not match the theoretical spectra used in standard identification algorithms. We resolve this difficulty using a modified search algorithm we call Conditional Ion Distribution Search (CIDS). Increased identifications from CIDS along with direct measurement of amino acid enrichment and statistical modeling that accounts for heterogeneous information across peptides, dramatically improves the accuracy and precision of half-life estimates. We benchmark the approach in cells, where near-complete labeling is possible, and conduct an in vivo experiment revealing, for the first time, differences in protein turnover between mice and naked mole-rats commensurate with their disparate longevity.

## Main

Protein turnover rates describe cellular phenomena that play important roles in the prevention of protein aggregation, damage reduction and preservation of proteostasis^1–3^. Mass spectrometry-based proteomics techniques have enabled the measurement of proteome-wide turnover rates *in vivo*^4^. However, these approaches present challenges not encountered when using techniques for cell cultures, such as Stable Isotope Labeling by Amino acids in Cell culture (SILAC)^5^. In a SILAC experiment, adding isotopic labels to the media quickly leads to the near complete replacement of free-floating amino acids with labeled counterparts. Similar procedures *in vivo* result in only partial labeling^6^ and the time until enrichment saturation varies across environmental conditions.

Data analysis typically assumes stable amino acid enrichment and is complicated by dynamic enrichment in both the beginning of an experiment as amino acids are being replaced and at the end of an experiment as degradation-driven recycling of amino acids alters enrichment levels^7^. Furthermore, enrichment saturation at relatively low levels has a negative impact on quantitative performance. This motivates the administration of high doses of labeled amino acids, but even a flooding dose may achieve only ~30% enrichment on a short time scale^6^ and the cost of isotopic labels for larger organisms can be substantial.

Deuterium oxide (D_2_O) can be used as an inexpensive alternative labeling strategy that results in the partial labeling of many amino acids^8^. However, in addition to the above challenges, identifying deuterated peptides presents unique challenges. Deuterated peptides often have isotopic distributions spread across many masses and the location of the largest peak depends on the amount of protein turnover that has occurred. Standard proteomics identification algorithms^9^ and modern machine learning models^10,11^ have been designed and optimized to identify monoisotopic peptides. When a peptide is isolated with an unknown number of heavy amino acids, two problems occur. First, the precursor mass may not match the masses created during in-silico digestion. Second, the resultant fragmentation spectra present non-linear shifts in fragment ion masses. Consequently, protein turnover experiments based on ubiquitous labeling strategies^12–15^ inevitably result in vanishing numbers of successful peptide identifications as the amount of protein turnover increases.

In this paper, we describe an experimental and mathematical framework for estimating protein turnover based on measured dynamic amino acid enrichment. From this framework, we derive an algorithm for identifying peptides that contain an unknown number of heavy isotopes. This approach effectively solves the problem of identifying highly deuterated peptides resulting in larger and more information rich datasets. Our methodology is then validated in cell culture by comparing our turnover rate estimates with the gold standard in the field, SILAC. Finally, we apply our approach to an in vivo study comparing mice and naked mole rats. Loss of proteostasis is one of the hallmarks of aging^3^ and these two species have widely divergent longevity, so interspecific differences in turnover rates may illuminate important features of the aging process.

## Results

### A framework for dynamic amino acid enrichment

Our objective is to estimate protein turnover rates from observed proportions of peptide isotopes. As outlined in the principles of Mass Isotopomer Distribution Analysis,^16,17^ determining the amount of turnover that has occurred requires knowing an initial and a final isotopic distribution for each peptide. The initial distribution can be derived from known natural isotopic proportions of each element. However, the final distribution, which we expect to see once a protein has been completely replaced with copies generated after the introduction of isotopic labels, typically requires the assumption of stable enrichment levels for each amino acid. When the enrichment levels are dynamic, this assumption is violated and the probability that the protein synthesis machinery will have selected a labeled amino acid becomes dependent on time.

Using metabolomics to observe the uptake of deuterium into each free-floating amino acid, we estimate the isotopic amino acid distributions for each time interval on which proteomics samples are analyzed (Figure 1a). We then estimate the marginal distribution of amino acid enrichment at each time interval and use these to calculate the distributions for each peptide synthesized during the interval. Thus, for each peptide at a given time we obtain initial and final state distributions determined from metabolomics and obtain one observed peptide distribution from proteomics. This framework for monitoring amino acid enrichment, predicting peptide isotopic distributions, and estimating protein turnover is the foundation for all of the advances presented in this paper (Figure 1b).

**Figure 1:**
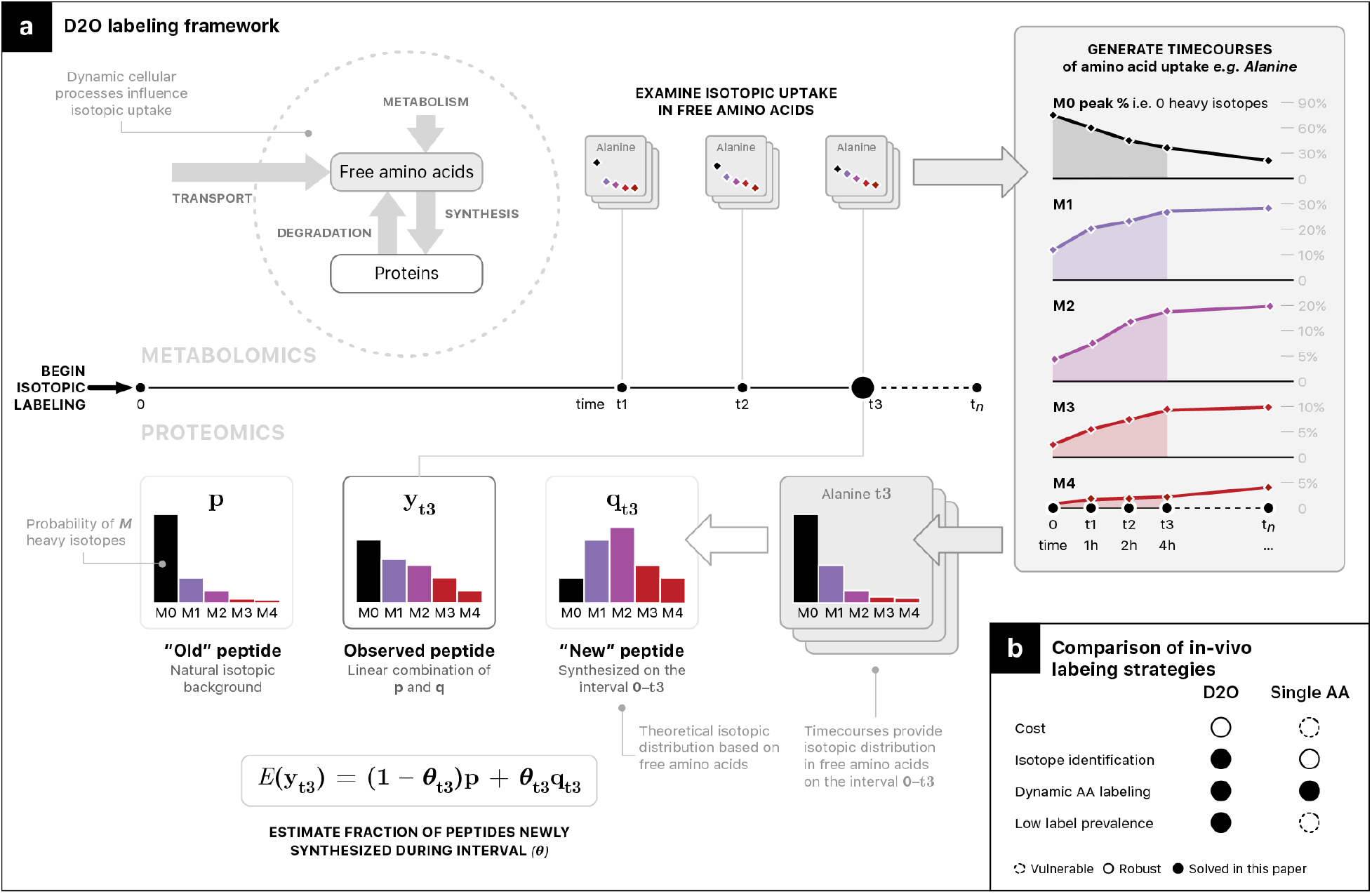
Experimental Framework for Turnover Estimation with Dynamic Enrichment. **a,** Experimental design. Isotopic envelopes are measured with both metabolomics (free-floating amino acids) and proteomics (peptides). Observing the uptake of amino acid labeling through time allows us to calculate the probability that a randomly selected amino acid, over a specific time frame, would have contained a heavy isotope. We use these probabilities to determine the theoretical isotopic distribution of a peptide synthesized during that interval. In this way we avoid assumptions about stable amino acid labeling, degradation and extracellular transport. **b,** A visual summary the problems solved with our strategy.

In theory, absent any error, the isotopic proportions observed in the proteomics experiment should be a convex combination of the old and new distributions. Accordingly, we create a Bayesian Dynamic Enrichment Model (BDEM) by treating observations as random variables from a Dirichlet distribution centered around the convex combination of isotopic initial and final states. Separate final states exist at each time point (the final state implies complete protein turnover, not the completion of sampling times) and are treated as constant in BDEM. The change in the percentage of proteins that existed prior to labeling is assumed to follow an exponential decay.

To inform both experimental design and data analysis, we consider two potential problems with our framework, both of which are intuitively obvious when the extreme cases are considered. If the old and new distributions are identical (a labeled peptide prevalence of zero) then turnover cannot be estimated. The consequences of low but non-zero labeling prevalence are less obvious. A second failure mode occurs if the concordance between turnover rates and sampling times is poor. At the extreme, if a protein has been completely turned over before the first sampling time or has not been turned over at all by the last time point, the ability to pinpoint the half-life with any level of precision is lost. We achieve a more general understanding of how concordance and prevalence impact quantitative performance through simulation studies.

### Label Prevalence and Sampling Time Concordance

We perform two simulations to evaluate the factors that impact estimation accuracy. Both studies simulate observations from a Dirichlet distribution with precision parameters and amino acid distributions taken from our validation experiment (Figure 2a). Labeled peptide prevalence manifests visually as a gap between the distributions of old and new peptides. Consider two tryptic peptides from the human mTOR protein, FDAHLAQAENLQALFVALNDQVFEIR and FDQVCQWVLK (Figure 2b). The larger peptide has a labeling prevalence of 86% while the smaller peptide has a prevalence of just 17%. Simulating random values centered at 70% between the boundaries, the difficulty in estimating turnover becomes visually apparent (Figure 2b). To quantify the impact of labeled peptide prevalence, we create an artificial situation where all of our simulated observations come from only a single level of prevalence.

**Figure 2:**
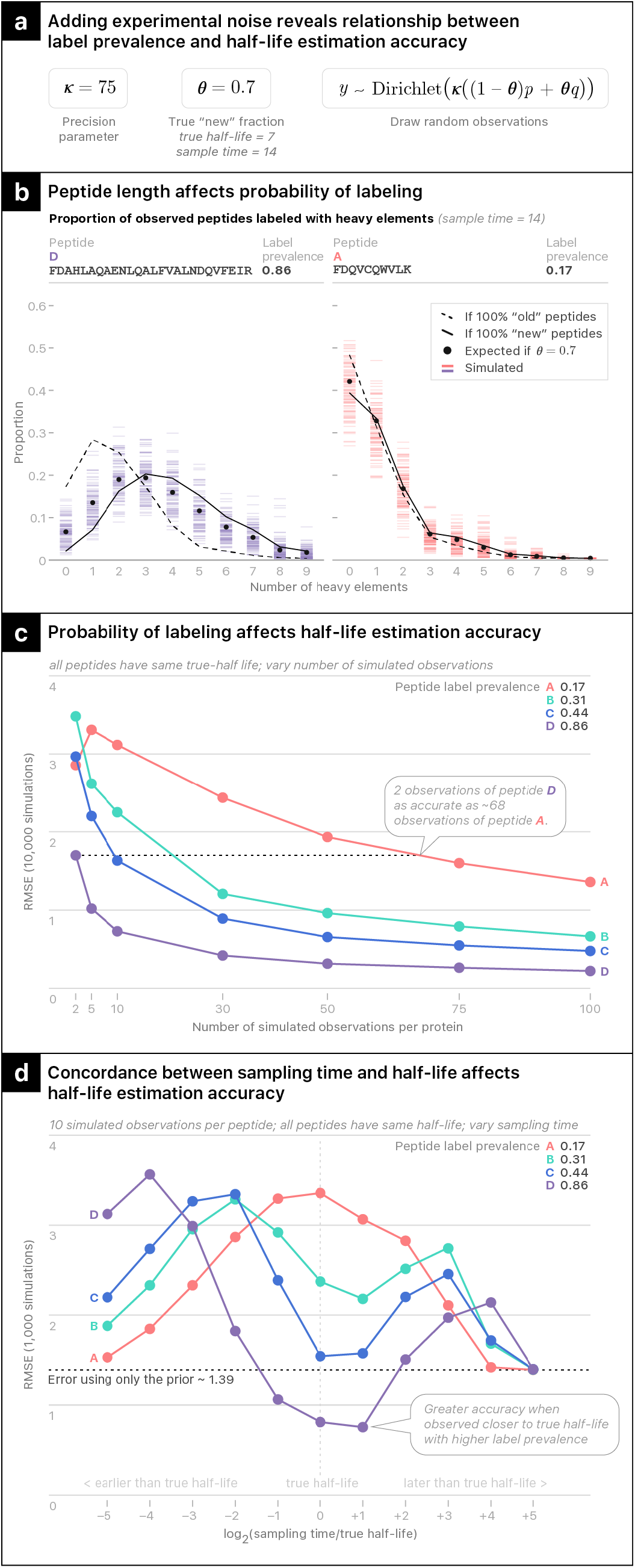
Peptide Labeling Prevalence and Sampling Time Concordance Largely Determine Quantitative Performance. **a,** The basic parameters of our simulation study including precision and amino acid enrichment were taken from the SILAC validation data. **b,** 100 simulated observations from small and large mTOR tryptic peptides with equivalent experimental error. Looking at the observations from the large peptide, six or seven isotopologues suggest that 70% of the protein had been turned over from visual inspection alone. For the small peptide, only the M0 peak provides much information. **c,** Peptide isotopes were simulated from the BDEM model with varying numbers of observations from a single peptide sequence for each protein. Half-life root mean squared error from 10,000 simulated estimates is plotted on the y-axis. Following the dashed horizontal line shows that two peptides with a label prevalence of 86% provide as much accuracy as approximately 68 observations with a prevalence of 17%. **d**, The sampling time was varied as two-fold multiples of the true half-life. As the sampling time deviates from the true half-life, in either direction, error converges to the error obtained when using the median of our prior as an estimate.

In the first simulation we vary sample size while keeping the true half-life of 7 days and the sampling time of 14 days fixed. After simulating 10,000 artificial proteins, we used the same model to estimate half-lives and plot the root mean squared error (RMSE; Figure 2c). Two observations from the peptide with prevalence of 86% provides the same level of accuracy as approximately 68 observations from peptides with a prevalence of 17%, underscoring the importance of peptide labeling prevalence.

In the second simulation, we fix the sample size at 10 observations but vary the simulated sampling times as two-fold multiples of the true half-life. Plotting the RMSE from this simulation (Figure 2d) we see that error generally remains lowest within a two-fold differential of the true half-life, with errors converging to what we would obtain using only our prior half-life distribution as the concordance decreases. This convergence occurs more quickly for low information peptides, resulting in an inverted curve for the peptide with a prevalence of 17%. This suggests that our Bayesian framework recognizes the low information content of peptides sampled far away from their true half-lives. In real experiments, samples will be collected at many time points, increasing the chances that at least one sampling time will be similar to the true half-life. Consequently, a well dispersed time course should provide better quantitative performance, but it will also result in heteroskedastic observations through time.

The peptides observed in a discovery proteomics experiment are not controlled. Consequently, none of the factors in these simulation studies (sample size, labeling prevalence, sampling time concordance) will be fixed across observations. The Bayesian BDEM automatically accounts for the heteroskedasticity (see Methods) but no statistical model will perform well if we fail to observe any high information peptides. Unfortunately, these are precisely the peptides that cause standard identification algorithms to fail.

### Identification of Peptides with Heavy Isotopes of Unknown Location and Quantity

Without a priori knowledge of the mass shift caused by the incorporation of isotopically labeled amino acids, standard identification algorithms fail for two reasons. First, the anticipated monoisotopic precursor mass will not match the observed *m/z*. Second, the process of mass isolating a population of ions that exclusively contain a fixed number of heavy isotopes, fundamentally alters the isotopic distribution of the fragment ions (Figure 3a). This isolation results in a non-linear shift along the *m/z* axis of the observed fragment spectra.

**Figure 3:**
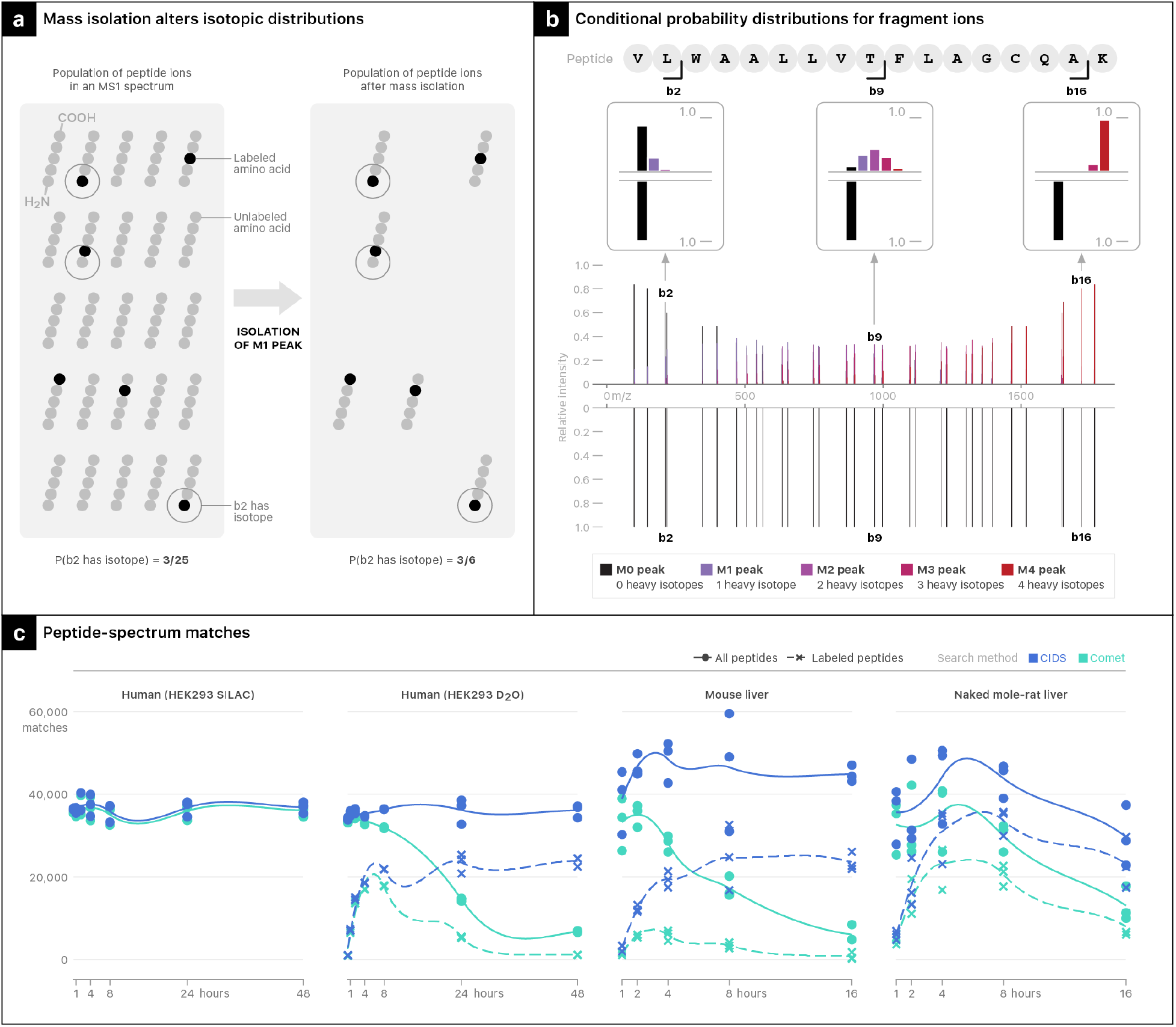
A New Algorithm for Identifying Labeled Peptides. **a,** Mass isolation alters the isotopic distributions fragment ions. **b,** Conditional probability distributions for each fragment ion are used to create a new theoretical MS2 spectrum (top) to replace the standard Comet spectrum (bottom). **c,** Loess curves through the number of peptide-spectrum matches at a protein FDR of 1%.

The first problem of a misaligned precursor mass can be trivially corrected by expanding the number of plausible masses that we search for. The second problem of the non-linear shift is more complex, but the behavior can be described according to the theory of conditional probability distributions (See Methods). For each potential peptide-spectrum match (PSM), we know the number of heavy isotopes that the precursor must contain within the mass tolerance of the instrument. Consequently, we can generate conditional probability distributions for each *b-* and *y*-ion given the candidate peptide and its number of heavy isotopes. Note that while this paper presents an algorithm for identifying peptides with the *b-* and *y-* ions, the theory could be extended to other fragmentation schemes. Substituting the isotopic distributions in place of the usual theoretical *b-* and *y-* ion peaks creates alternative theoretical spectra to use for peptide-spectrum matching (Figure 3b). We incorporated this approach into our installation of the open-source identification package Comet^18^.

In 51 deuterated samples collected from three species (mice, naked-mole rats, and human HEK293 cells) we consistently see the same pattern of the number of PSMs decreasing with time (Figure 3c). However, when using our Conditional Ion Distribution Search (CIDS), the number of PSM’s at the end of each experiment are similar to counts observed before much turnover occurred.

The gain in PSMs is only a small part of the advantage provided by CIDS. By counting only peptides with a labeling prevalence greater than 50% we can see that it is precisely the peptides that are most likely to be labeled that our algorithm recovers (Figure 3c). The algorithm not only increases the number of observations, it also recovers a set of peptides that are fundamentally more valuable than the ones identified with a standard search algorithm.

It should be mentioned that the theory underlying CIDS is not exclusive to isotope labeling protein turnover experiments. When a monoisotopic peak is isolated, the algorithm will produce an MS2 identical to the standard approach. This is why the PSM counts for the SILAC turnover data are only slightly increased from the standard Comet search (Figure 3c). For the vast majority of scans, the MS2 spectrum remains unaltered.

### Methodological Comparison with SILAC

In order to validate the theoretical advances described above, we analyzed protein turnover in HEK293 cells using both SILAC and D_2_O and applied our novel methodologies to these data. When the media can be controlled, the SILAC approach will result in peptides that are consistently labeled at nearly 100% prevalence on every peptide. In this sense, the SILAC measurements are optimal and serve as a useful ground truth for evaluating the quantitative performance of our D_2_O methodology.

We compared turnover from cells in media containing 100% heavy lysine, to the same cell line with the media replaced by 8% D_2_O, a commonly targeted concentration in blood which is limited by toxicity concerns (Figure 4a). Three replicates were collected at 0.5, 1, 2, 4, 8, 24 and 48 hours. The D_2_O data were searched with both Comet and CIDS and the resultant datasets were analyzed with BDEM and two previously described software packages used routinely to analyze D_2_O data, Deuterator^19^ and d2ome^20^. Each of these approaches removes outliers caused by errant peaks. Consequently, the number of uniquely quantified protein half-lives varies substantially across methodologies (Figure 4b). Filtering algorithms from d2ome and Deuterator are far stricter than our own requirements. Accordingly, the BDEM+CIDS approach results in 3,967 uniquely quantified proteins which represents a 99% increase over Deuterator+Comet (1,991 proteins) and a 209% increase over d2ome+Comet (1,285 proteins). However, the BDEM+CIDS approach still quantifies fewer proteins than the SILAC methodology (3,967 to 4,507) which did not utilize any quality control filtering.

**Figure 4:**
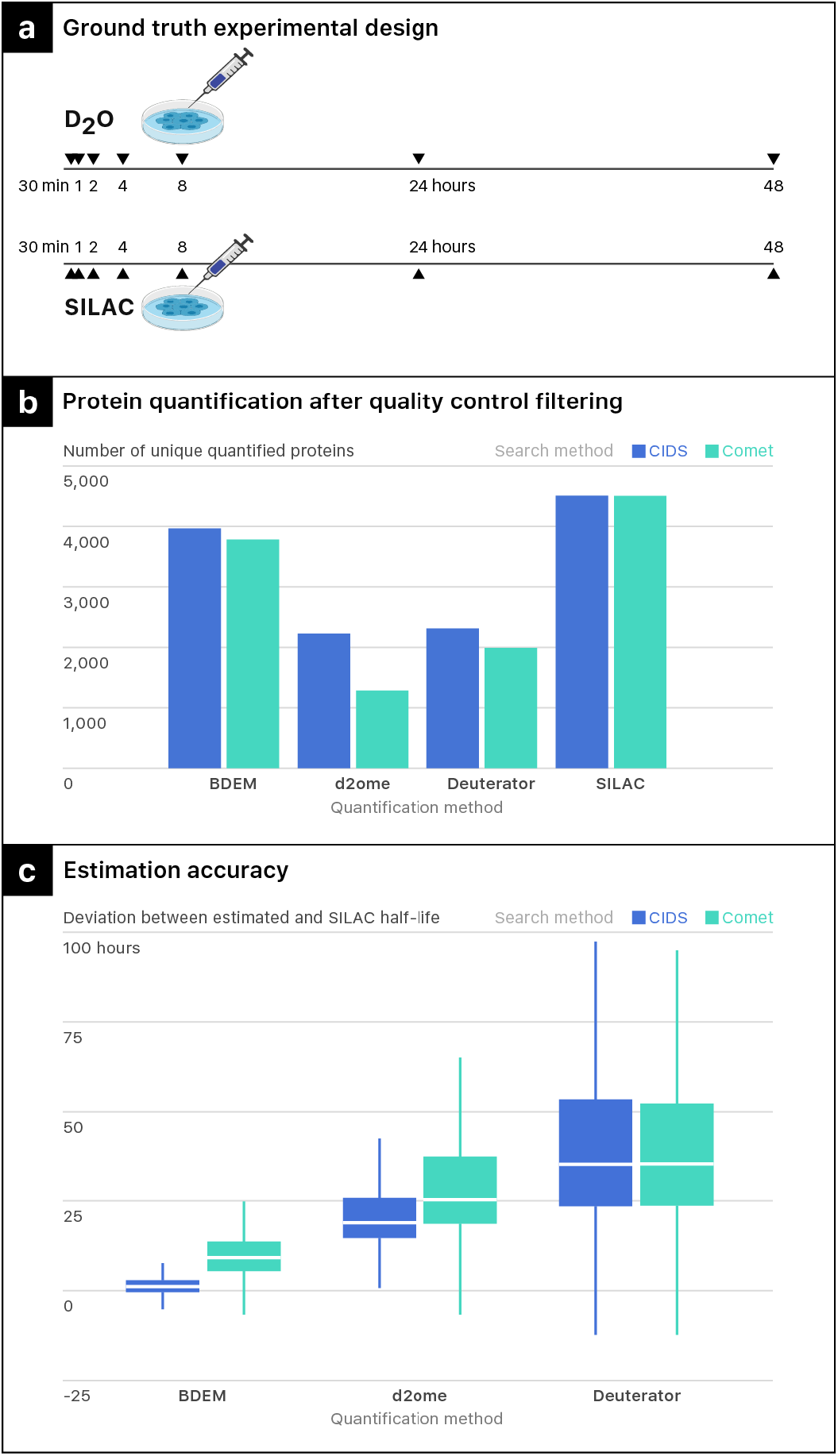
Evaluating Performance with a SILAC Ground Truth Experiment. **a.** Design of a ground truth experiment. In a HEK293 cell culture we compare SILAC labeling (100% replacement of lysine with a heavy isotope) against D2O labeling (8% D2O in the media). **b.** Counts of uniquely quantified proteins across quantification and identification algorithms. Data are comprised of three replicates at each timepoint (4, 8, 24 and 48hrs). All half-life estimation algorithms have filtering criteria that result in different total counts even on the same data. **c.** Boxplot of errors. Using the SILAC turnover data as ground truth, we show the deviation across algorithms. Error was calculated for each protein quantified by all of the algorithms (N=957).

Although CIDS alone provided only modest gains to the count of uniquely estimated protein half-lives, the effect on deviation from the SILAC results were profound. The full set of overlapping proteins across methods shows a loss of both precision and accuracy when using the standard Comet search, with a median absolute error of 9.17 hours. In contrast, the median absolute error using CIDS is only 1.18 hours (Figure 4c). The gains in accuracy of BDEM+CIDS compared with d2ome (median absolute errors of 18.92 and 25.33 hours for CIDS and Comet respectively) and Deuterator (median absolute errors of 35.39 and 35.48 hours for CIDS and Comet respectively) are greater still, and strongly suggest that the principles outlined in this manuscript are essential for accurate half-life estimation.

### Slower Protein Turnover in Naked Mole-Rats Compared to Mice

We next applied our new methodologies to evaluate interspecific differences in protein turnover in the long-lived (~37y), cancer resistant naked mole-rat and the similar-sized, cancer prone, short-lived (~4y) C57BL/6 mouse^21^. Unlike laboratory mice raised on synthetic diets, captive naked mole-rats are fed a low protein, vegetable-based diet mimicking their natural diet. Moreover, as in the wild, captive naked mole-rats obtain water entirely from their food and are not provided drinking water, making standard D_2_O labeling strategies impossible. To circumvent these challenges, we administered daily intraperitoneal (IP) injections of D_2_O for 16 days and periodically collected liver samples for metabolomic and proteomic assessments (Figure 5a). C57BL/6 mice were subjected to the same protocol. The IP administration successfully labeled free floating amino acids in the liver, revealing both consistent patterns in protein turnover rates across species and exceptions that coincide well with the literature on naked mole rat biology.

**Figure 5:**
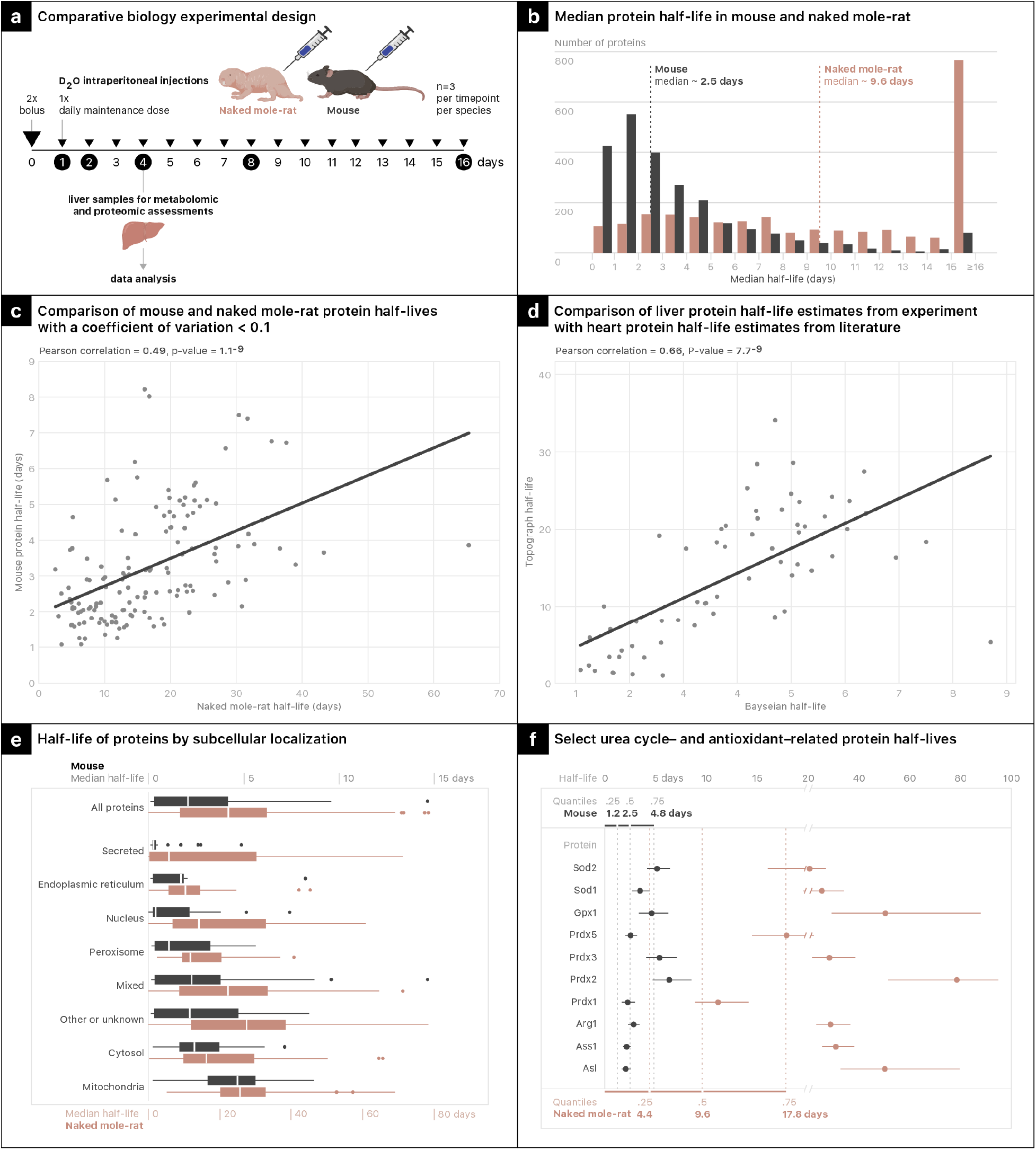
Comparative Biology of Protein Turnover in Mice and Naked Mole Rats. **a.** Design of a comparative biology experiment. **b.** Distribution of half-life estimates for each species. All quantified proteins are included. **c.** Scatterplot and correlation of precisely (CV < 0.1) estimated half-lives between species. **d.**Scatterplot between half-life estimates from our mouse liver data compared with previously published turnover data from young mouse hearts. Only proteins with a reported coefficient of variation less than 0.1 are shown. **e.** Subcellular localization of precise half-lives. **f.** 99% posterior half-life distributions for select proteins.

Histograms of protein half-lives in the two species show a systemic shift in the turnover rates, with a median half-life of 2.5 days in mice (3,547 proteins quantified) and 9.6 days in naked mole-rats (3,200 proteins quantified, Figure 5b). This relationship was only slightly changed by restricting our analysis to highly precise half-life measurements with coefficients of variation less than 10%. Slower turnover in naked mole-rats when compared to mice is consistent with published studies conducted in cultured cells from these species using a pulse-SILAC approach^2^. Despite the substantial overall difference in turnover rates, we still see a significant correlation between half-lives across species (Figure 5c). Moreover, in spite of different analysis platforms, labeling strategies and even tissues being analyzed, we see a slightly higher correlation when comparing our mouse liver turnover rates to previously published findings on turnover in the heart^22^ (Figure 5d). Consistent interspecies patterns are also seen when grouping rates by subcellular localization, with slower rates in the mitochondria and faster turnover in the nucleus and endoplasmic reticulum (Figure 5e). These patterns provide further validation of our methodology while also enabling the identification of proteins that defy these typical relationships.

Many proteins with average turnover rates in mice are among the slowest to turn over in naked mole rats (Figure 5f). All three cytosolic urea cycle enzymes (argininosuccinate lyase [ASL], argininosuccinate synthase [ASS1] and arginase [ARG1]) are more stable than we would expect based on the results in mice. Living in sealed underground burrows with designated latrines, naked mole-rat colonies commonly encounter gaseous atmospheres low in oxygen and high in both carbon dioxide and ammonia^23^. The animals are exceedingly tolerant of such hostile conditions and, strikingly, do not avoid ammonia-saturated atmospheres. Since the urea cycle detoxifies ammonia, naked mole-rats likely rely heavily on this pathway.

In addition to the urea cycle, the turnover of many proteins implicated in the breakdown of harmful reactive oxygen species, such as peroxiredoxins (PRDX), glutathione peroxidase (GPX1) and superoxide dismutases (SOD), are markedly slower than expected (Figure 5f). Response to oxidative damage is an active area of naked mole-rat research with previous studies reporting enhanced cytoprotection in response to oxidative damage through upregulation of the cytoprotective molecule Nuclear Factor Erythroid 2-related Factor 2 (NRF2) and concomitant upregulation of antioxidants, detoxicants and molecular chaperones^23^.

## Discussion (No Subheadings)

Our proposed methodology greatly increased the depth of discovery and the precision of turnover rate estimation. These gains were validated *in vitro* and *in vivo*, highlighting the importance of five fundamental properties of protein turnover experiments.

First, quantitative performance depends on labeled peptide prevalence, which can be altered either by increasing amino acid enrichment, or by selecting peptides with increased opportunities for enrichment. The prevalence-peptide selection dynamic should be considered when selecting labeling strategies and digestion agents. Understanding the quantitative value of high label prevalence should prove especially useful when applying targeted proteomics techniques to prespecify peptides of interest^24,25^.

Second, in the presence of dynamic amino acid enrichment, the probabilities relevant to protein turnover vary with time. These probabilities can be estimated by monitoring the enrichment of free-floating amino acids in the tissue of interest. Dynamic enrichment resulting from either inconsistent label administration or amino acid recycling has long been a concern even in the context of bulk synthesis measurements. The dynamic enrichment model presented here addresses this challenge.

Third, the concordance between sampling times and true half-lives alters estimation precision, resulting in nothing more than reliable lower or upper bounds when the deviation becomes extreme. Yet these bounds remain highly informative, requiring careful assessment of uncertainty intervals whenever the concordance is low.

Fourth, in a mass spectrometry experiment, the mass isolation of an unknown number of heavy isotopes alters the isotopic distribution of each fragment ion. Utilizing conditional probability distributions to describe this behavior enables the collection of the peptides that were most likely to have been labeled. While D_2_O labeling motivated the creation of the CIDS algorithm, any mass spectrometry experiment that isolates ions with an unknown number of heavy isotopes could benefit from the approach.

Finally, heteroskedastic observations should be anticipated in protein turnover studies. While some labeling approaches may keep labeled peptide prevalence constant, no methodology can keep sampling time concordance fixed through time. Statistical modeling that accounts for these factors should be used to minimize estimation error.

Despite the seemingly noisy observations, D_2_O can be used to reliably study protein turnover. Advances in the quantitative performance of D_2_O turnover experiments are especially valuable considering the cost differential between various isotopic labels. At the time of this writing we estimate the cost of labeling with a heavy isotope of leucine^22^ to be approximately 35 times more expensive than our D_2_O labeling strategy. However, the above principles offer more than cost savings, as many concepts apply to protein turnover experiments regardless of the labeling strategy. In particular, labeled peptide prevalence and sampling time concordance should always be taken into account when designing experiments and creating statistical models.

Taken together, the present advances have dramatically enhanced our capabilities for studying proteome-wide turnover rates. We have effectively eliminated the requirement for stable amino acid enrichment while simultaneously describing a strategy for extracting meaningful signals even when overall enrichment levels remain low. This combination of developments opens numerous possibilities for exploring currently unseen aspects of *in vivo* biology.

## Methods

### Calculating Peptide Isotopic Distributions

We now consider the problem of deriving isotopic distributions both from the perspective of a ribosome randomly retrieving individual amino acids from the free-floating pool of available amino acids.

Let *A*_*it*_ be a random variable denoting the number of heavy isotopes contained in an amino acid selected by the ribosome during the interval (0, *t*), where, *i* = 1, ..., 20, indexes the standard DNA encoded amino acids. *A*_*it*_ is a discrete random variable mapping to a subset of the non-negative integers. For the D_2_O labeling discussed in this paper the range is typically confined to the set [0, 1, 2, 3], but other labeling strategies could look substantially different. Note, that in forcing the range to take non-negative integers we are ignoring small differences in isotopic masses. For example, an amino acid containing a single heavy hydrogen and the same amino acid containing a C13 isotope have distinct masses but, in our framework, both amino acids take a value of *A*_*it*_ = 1. The ability to distinguish these masses in a mass spectrometer is diminished for larger analytes, but in our framework, we ignore the discrepancy throughout the entire mass range.

The probability that the *i*th amino acid, selected by the ribosome between (0, *t*), contains *x* heavy isotopes can be written as *p*(*A*_*it*_ = *x*) and we refer to the function defining these probabilities for all x as 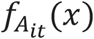 If the set of amino acid distributions, 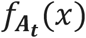 is known, either through metabolomics measurements or from a priori knowledge, and if the incorporation of heavy labeled amino acids is mutually independent for all amino acids in the sequence, then the isotopic distribution of a peptide can be modeled as the distribution of a sum of discrete random variables.

Let *H*_*n*_ be a random variable denoting the number of heavy isotopes contained in peptide *n*, *n* = 1, ..., *N*, where *N* represents the total number of unique peptides observed in an experiment. If peptide *n* consists of M amino acids from the set [*A*_*it*_, ..., *A*_20*t*_] (drawn with replacement), then the isotopic distribution of the peptide is 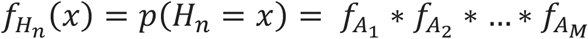, where * denotes the discrete convolution operation. In this way, we can quickly derive the isotopic distribution of a randomly selected peptide once we know the isotopic distributions of the constituent amino acids. This formula is not exactly correct, as the process of binding two amino acids involves the loss of 2 hydrogens and 1 oxygen, each of which may have contained heavy isotopes. Accordingly, we should deconvolve the natural isotopic distribution of H_2_O, M-1 times from the peptide isotopic distribution.

When using the natural isotopic proportions of each amino acid the convolution operation provides the isotopic distribution of a peptide prior to any experimental perturbation. We will refer to this as the “old peptide distribution” and use the shorthand notation for peptide *n*,

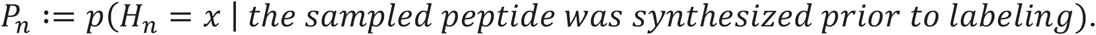

For peptides that were synthesized after *t* = 0, we rely upon direct observation of the amino acid enrichment through time to estimate each probability. In this scenario the amino acid probabilities may change through time, but we will still be able to estimate the probability that an amino acid, selected during a specific time interval by the ribosome, contains x heavy isotopes. In this case we use the shorthand

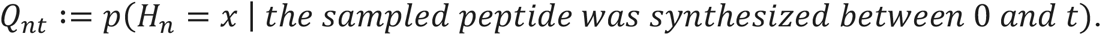

Neither *P*_*n*_ or *Q*_*nt*_ define the isotopic distribution of a peptide sampled at time *t*. Rather, they represent complementary subpopulations. To find the peptide population distribution at time *t* we need to know the proportion of newly synthesized peptides. Let the percentage of newly synthesized copies of peptide *n* at time *t* be denoted as *θ*_*nt*_ and for each sampled peptide let *L*_*n*_ = *o* if peptide was synthesized prior to labeling (“o” is for old) and *L*_*n*_ = *e*, (“e” is for new) if the peptide was synthesized between (0, *t*). Then the isotopic distribution of peptide *i* at time *t* is given by

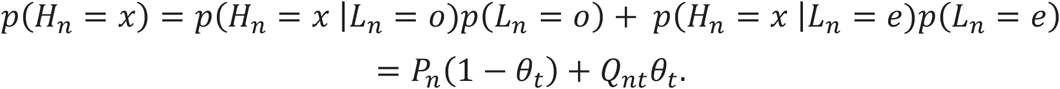

### Deriving the Isotopic Distributions of the Fragment Ions Used for Peptide Identification

Mass spectrometers collect ions and present a spectrum of mass-to-charge (*m/z*) ratios on one axis, and intensities (more precisely, ion counts, measured in arbitrary intensity units) on the other axis. This readout of many masses is often referred to as an MS1 spectrum since it is the first in a sequence of scans. Because peptide identifications cannot be uniquely determined by a mass alone, mass spectrometers will select a peak from the MS1 scan, isolate ions close to the corresponding mass and fragment them. The mass spectrum of fragment ions (MS2) will then be compared against theoretical fragmentation spectra from all of the peptides that are consistent with the mass of the isolated precursor^9,26^. The most common approach for fragmenting peptides results in series of *b-* and *y-* ions^26^. Our goal is to create conditional isotopic distributions for each b- and y-ion from a candidate peptide. For simplicity, we assume a mass isolation window sufficiently narrow to select only precursor ions from one isotopologue. (often only true for precursors with a charge state of 2 since a standard isolation window has a width of 1 Da). If the monoisotopic peak was selected, then no amino acids in the isolated set of peptide ions contain any heavy isotopes. Likewise, the *b* and *y* fragment ions will not contain any heavy isotopes. This is also true for labeled monoisotopic peaks. For example, in a SILAC experiment we might isolate a peptide with isotopically labeled lysine and all fragment ions will contain exactly one mass for lysine at a predictable offset. The fragment ion peak masses only become difficult to predict when we isolate a precursor containing a heavy isotope at an unknown position. For example, if we isolate ions containing a single ^13^C isotope, that isotope could have been present in any of the amino acids within the peptide. The whole peptide must contain exactly one heavy amino acid, but the location of the heavy amino acid will change throughout the population of ions. Consequently, each *b-* and *y*-fragment ion will have a predictable distribution of isotopes that can be helpful in identifying the precursor.

Let *H* represent the number of heavy isotopes contained in an isolated precursor peptide. For each candidate peptide, we know ℎ, since only one possibility will be within the mass tolerance of the instrument. We let *θ* represent the percentage of peptides that, at the time of sampling, had been synthesized after label administration. Marginal isotopic distributions of *b*- and *y*-ions (the proportions we would expect to see if all peptides were fragmented) can be generated using the concepts described for calculating peptide isotope distributions. For a peptide of length *M*, let *B*_*i*_ and *Y*_*j*_ represent random variables for the number of heavy isotopes contained in randomly sampled *b*_*i*_ and *y*_*j*_ ions respectively (*i* = 1, ..., *M* and *j* = 1, ..., *M*). Further, let 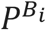,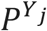,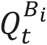 and 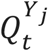 represent a shorthand for the marginal isotopic probability distributions of each *b* and *y* ion where, as before, *P* denotes a distribution prior to labeling and *Q*_*t*_ represents the isotopic distribution of fragments synthesized between (0, *t*). These are the distributions of the respective populations as they would be seen in a cell, without restricting the total number of heavy isotopes found in the precursor. We present the derivations for the b ions (they are analogous for the y ions). Mass isolation results in the following conditional probability distribution:

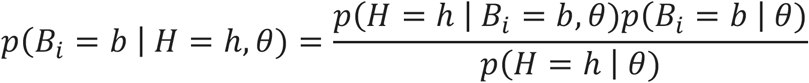

Both *p*(*B*_*i*_ = *b* | *θ*) and *p*(*H* = ℎ | *θ*) can be calculated using the standard convolutions previously described. Calculating *p*(*H* = ℎ | *B*_i_ = *b*, *θ*) requires once again separating out old and new peptides. If *L* defines the old versus new status of the peptide, as before, then we have

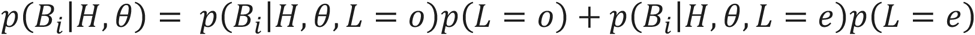

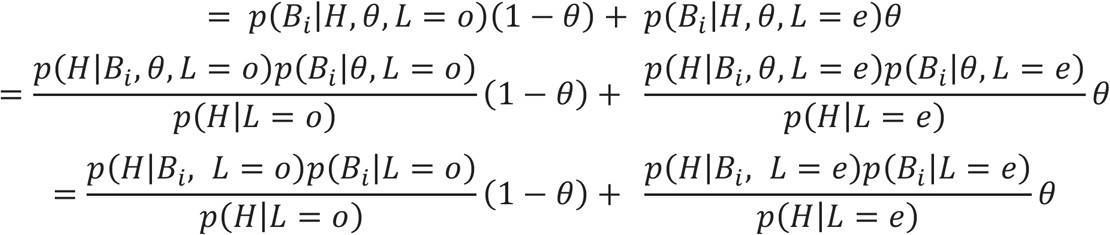

Each of the probabilities in this equation can be found using convolutions of natural and observed amino acid sequences.

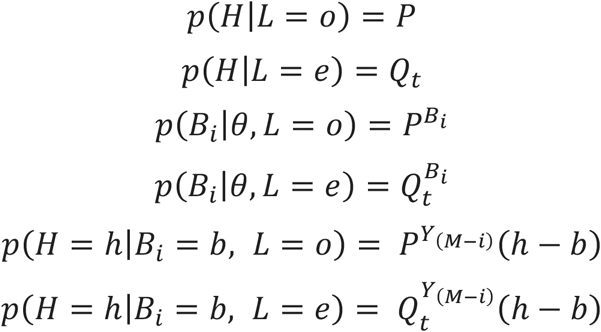

The last two expressions follow from the complementary nature of *b* and *y* ions and the observation that

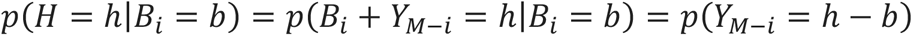

A shorthand analytic expression for the conditional distribution of a *b* ion (as will be observed in an MS2 spectra) is given by

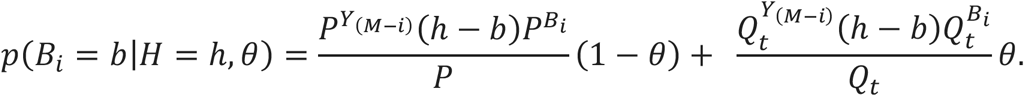

This expression is still dependent on the unknown parameter *θ*. A simple solution to this problem is to treat *θ* as a uniform random variable and to integrate it out of our distribution.

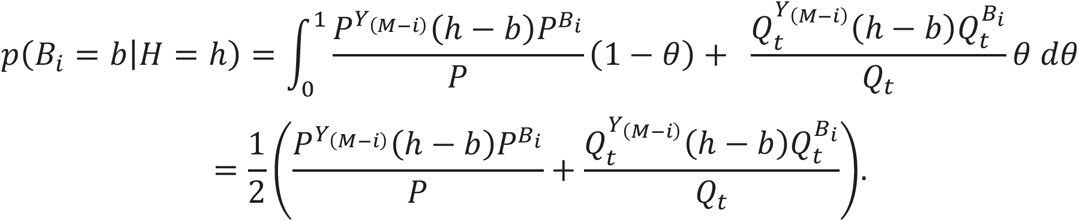

From these equations we are able to generate alternative theoretical spectra for use in a proteomics search algorithm. Placing these distributions at their corresponding masses results in a new theoretical MS2 spectrum (See Figure 3b).

We modified the open source identification package Comet^18^ to allow for conditional isotope distribution searches (CIDS). An R script for generating the MS2 distributions has been included in the supplemental files.

All searches performed in this manuscript follow a standard identification workflow using a reverse hit database^27^. Linear discriminant analysis was used to reduce the protein level false discovery rate to 1%^28^.

When searching the SILAC data, the above equations are greatly simplified since there is only one relevant set of amino acid isotopic distributions. However, the benefits are also minimal which is to be expected since the vast majority of MS2 scans will result from isolating a monoisotopic peak. We have also observed that averaging the old and new fragment ion distributions only results in minor improvement beyond what we obtain when using only a single set of amino acid distributions. This suggests that, conditional on the number of isotopes in the precursor, the amount of turnover has a relatively minor impact on the isotopic distributions.

### Statistical Modeling of Protein Turnover

For a unique protein *w*, (*w* = 1, ..., *W*), suppose that we observe *N*_*wtk*_ peptides at time t, (*t* = 1, ..., *T*) in replicate *k*, (*k* = 1, ..., *K*). Further assume that for each observation ***y***_***n***_, (*n* = 1, ..., *N*_*wtk*_), the underlying isotopic distributions, (***P***_*n*_, ***Q***_*n*1_, ..., ***Q***_*nt*_), are all known. In the rest of this section bold font will be used for vectors. Then if *θ*_*wt*_ denotes the percent of newly synthesized copies of protein *w* at time *t*, in the absence of any sampling or experimental error we would expect the isotopic proportions observed in a mass spectrometer to be given by ***P***_***n***_(1 − *θ*_*wt*_) + ***Q***_***nt***_*θ*_*wt*_.

Unfortunately, there will be both sampling variability and experimental errors that contribute to deviations from these expectations. For observations from a single protein, we define each measurement, ***y***_***ntk***_, as a random draw from a Dirichlet distribution centered around the convex combination of old and new distributions.

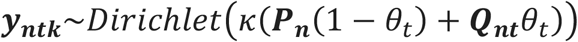

In this parameterization of a Dirichlet distribution, the precision parameter *κ* is a scalar multiplier of the proportion vector defined by the convex combination.

Further defining our data structure, we need to connect the percentage of newly synthesized proteins to an overall turnover rate. To this end we use the standard exponential decay model, using a half-life, η, to define the rate of protein turnover.

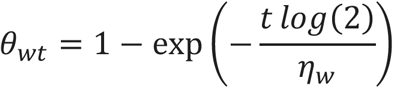

From this basic structure there are many possible approaches to estimation and inference. We chose to fit a Bayesian model which offers a convenient framework that synthesizes the complicated heteroskedastic data structure into a single posterior distribution describing our updated beliefs about a protein half-life. The full specification for a set of *M* proteins where peptide *n* is nested within each protein-time-sample combination (*m*, *t*, *k*), is given by

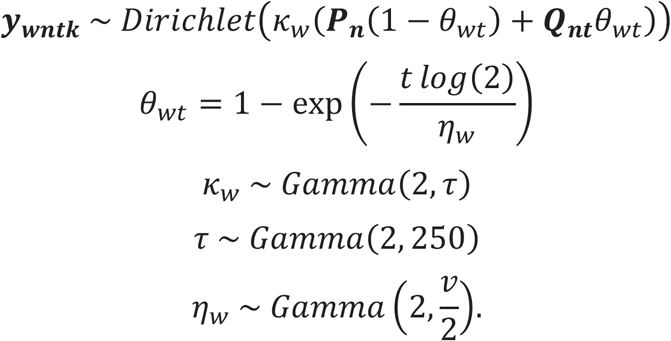

τ establishes an average precision throughout the dataset. Defining *k*_*w*_ as a random variable with mean 2τ results in a model that will “borrow” information about the precision of measurements when sample sizes are low. Similar strategies have been used for dealing with imbalanced data from relative abundance proteomics experiments^29^.

The three gamma distributions all use a shape and scale parameterization. The prior shape parameter of 2 was selected to avoid computational difficulties related to sampling near zero^30^.

We fit the above model in the Rstan package for modeling with the Stan programming language^31^. We report credible intervals taken as the 1^st^ and 99^th^ percentile of the posterior half-life distributions and we use the median of this distribution as a point estimate for each half-life. A few comments about the modeling and interpretation are necessary.

Recent applications of Bayesian modeling to mass spectrometry proteomics data make use of weakly informative and non-informative priors^29,32^. These priors are selected to make it highly unlikely that changes in the prior will have a noticeable effect on the final results, while simultaneously avoiding problems caused by sampling from unrealistic regions of the parameter space^33^. The Gamma(2, 250) distribution was chosen with the weakly informative principle in mind. Across many datasets, the average precision parameter always fell well within a Gamma(2, 250) distribution and given the large number of observations contributing to this parameter it is unlikely that reasonable changes to the prior would have a noticeable effect on the posterior.

In our software, the hyper-parameter *v*, which represents the mean of the half-life distribution, can be entered by the user. For each analysis in this paper, the parameter was set to 10, but this will not always be an appropriate choice. Unlike the other parameters, η_*w*_, has an informative prior. The mean should be selected to create a prior that genuinely reflects what the distribution of half-lives is believed to be. This belief should change depending on the sampling times of the experiment, the model organism and the tissue being analyzed.

The primary motivation for using an informative prior on the half-lives is the problem of sampling time concordance. When a true half-life is far removed from the last sampling time, the observations will be consistent with a range of half-lives that are unbounded in one direction. Suppose that we have an experiment with a final sampling time of 10 days, and we measure peptides from a protein with a true half-life of 100 days. Intuitively, the measurements will show that almost no turnover occurred. Such measurements are consistent with a half-life of 100 days, but they are also consistent with a half-life of 1,000 years. It is simply impossible to pin down the exact half-life with any level of precision. Although a precise point estimate for the turnover rate cannot be obtained, the data still contain valuable information regarding the lower bound of the rate. In our thought experiment, we can say with great confidence that the half-life is greater than 10 days. A more exact estimate of the lower bound will depend on both the sampling times and the measurement variability. The Bayesian model provides a relatively straightforward approach for establishing a lower bound, and an informative prior on the half-lives helps to avoid problems caused by sampling values from an unbounded domain.

Extra care is required when interpreting half-lives for proteins with poor sample time concordance. When a posterior half-life distribution is imprecise (we used a coefficient of variation (CV) > 10% as an arbitrary cutoff) and the median falls above our sampling range, we are only interested in the lower limit of the distribution. On the other side of the sampling range, if the posterior has a CV > 10% and the median falls below our first sampling time, then we only pay attention to the upper limit. In both cases, results must be interpreted with care since the prior will become increasingly influential as sampling time concordance decreases.

### Metabolomics

Cell culture samples were prepared for metabolomics by centrifugation at 14,000g for 5 min, followed by aspiration of the supernatant, followed by evaporation of the supernatant under nitrogen gas. Samples were resuspended in 200 μL of water by vortex mixing for analysis via the ion pairing method described below. A 40 μL aliquot of the resuspension was then diluted with 160 μL of acetonitrile, combined by vortex mixing, and centrifugation at 14,000g for 5 min.

Metabolites in the supernatant were analyzed in positive ionization mode using a Thermo Scientific QE-plus mass spectrometer coupled to a Thermo Scientific Vanquish UHPLC, Amino acids were separated using a SeQuant^®^ ZIC^®^-pHILIC column, 5μm particle size, 200Å, 150 × 2.1 mm. Mobile phase A was 20 mM ammonium carbonate in water (pH 9.2); mobile phase B was acetonitrile. The flow rate was 150 μL/min and the gradient was t = −6, 80% B; t= 0, 80% B; t= 2.5, 73% B; t=5, 65% B, t= 7.5, 57% B; t= 10, 50% B; t= 15, 35% B; t= 20; 20% B; t= 22, 15% B; t= 22.5, 80% B; t= 24; 80% B. The mass spectrometer was operated in positive ion mode using data-dependent acquisition (DDA) mode with the following parameters: resolution = 70,000, AGC Target = 3.00E+06, Maximum IT (ms) = 100, Scan Range = 70 to 1050. The MS2 parameters were as follows: resolution = 17,500, AGC Target = 1.00E+05, Maximum IT (ms) = 50, Loop Count = 6, Isolation Window (m/z) = 1, (N)CE = 20, 40, 80; Underfill Ratio = 1.00%, Apex Trigger(s) = 3 to 10, Dynamic Exclusion(s) = 25.

Liver samples were homogenized whole in 1.5mL microcentrifuge tubes in 750μL of −80℃ 80% methanol, 20% water with one 7mm stainless steel bead using a 2010 Geno/Grinder (SPEX) with a run time of 45 seconds at 1,750 strokes/minutes, followed by a 30 second rest, repeated twice. The homogenization protocol was repeated until samples were fully homogenized as determined by visual inspection, resting on dry ice between each cycle. Samples were then spun at 4℃, 8,000RCF to pellet proteins. Supernatant was processed for all amino acids except cysteine as follows: 300μL of supernatant was evaporated under nitrogen at 4℃ and resuspended in 600μL of 80% acetonitrile, 20% water for LC-MS/MS analysis. For determination of cysteine, cysteine was derivatized by combining 200μL of supernatant with 1.2mL of 80% methanol, 20% water, and 1.5mL of derivatization buffer consisting of 80% methanol, 20% water, 10mM sodium bicarbonate, pH’ed to 7.4 using formic acid, containing 100mg/mL N-Ethylmaleimide (NEM). Samples were incubated for four hours, evaporated under nitrogen at 4℃ and resuspended in 600μL of water for LC-MS/MS analysis.

All amino acids were analyzed on a Thermo Scientific QE-plus mass spectrometer coupled to a Thermo Scientific Vanquish UHPLC. All data were acquired in profile mode. Non-cysteine amino acids were separated on an Agilent Infinity Poroshell 120 HILIC-Z column (2.1mm x 100mm, 2.7 μm particle size) as previously described^34^ with the following modifications: Mobile Phase A was 20mM ammonium formate at pH 3. Mobile Phase B was 90% acetonitrile, 10% water, with 20mM ammonium formate at pH 3. Total run time was 15.7 minutes at 0.4mL/min with the following gradient, 100% mobile phase B was ramped to 70% over 11.5 minutes, the column was washed at 50% B for 1 minute, and re-equilibrated at 100% B for 3 minutes. Mass spectrometry data were acquired in positive ion mode using data-dependant acquisition (DDA) mode with an inclusion list. Full MS was acquired at 70,000 resolution with an AGC target of 5e5 with 50 ms maximum injection time. Data-dependent MS2 was collected at 17,500 resolution with an AGC target of 1e5, max fill time of 50 ms, and stepped collision energy of 20, 40, 80.

Samples were analyzed for NEM-derivatized cysteine using a reverse phase ion-pairing chromatographic method with an Agilent Extend C18 RRHD column, 1.8μm particle size, 80Å, 2.1 × 150 mm. Mobile phase A was 10 mM tributylamine, 15 mM acetic acid in 97:3 water:methanol pH 4.95; Mobile phase B was Methanol. The flow rate was 200 μL/min and the gradient was t=−4, 0% B; t=0, 0% B; t=5; 20% B; t=7.5, 20% B; t=13, 55% B; t=15, 95% B; t=18.5, 95% B; t=19, 0% B; t=22, 0% B. The mass spectrometer was operated in negative ion mode using data-dependent acquisition (DDA) mode with the following parameters: resolution = 70,000, AGC Target = 1.00E+06, Maximum IT (ms) = 100, Scan Range = 70 to 1050.The MS2 parameters were as follows: resolution = 17,500, AGC Target = 1.00E+05, Maximum IT (ms) = 50, Loop Count = 6, Isolation Window (m/z) = 1, (N)CE = 20, 50, 100; Underfill Ratio = 1.00%, Apex Trigger(s) = 3 to 12, Dynamic Exclusion(s) = 20.

Metabolites were identified by matching fragmentation spectra and retention times from chemical standards that were previously analyzed on the same instrumentation. Identity, isotopic peaks, and peak integration were manually verified and quantified using our modified version of MAVEN^35^.

### Proteomics

Water, HEPES, urea, organic solvents, Pierce Detergent Compatible Bradford Assay Kit and Pierce High pH Reversed-Phase Peptide Fractionation Kit were purchased from Thermo Fisher Scientific (Waltham, MA). Lys-C was purchased from WAKO Chemicals (Richmond, VA). Oasis HLB 96-well uElution Plate was purchased from Waters Corporation (Milford, MA). Unless otherwise stated, all other chemicals were purchased from Sigma.

Samples (liver, muscle, cells) were added to 80% methanol, 20% water (1 mL) and centrifuged (15 mins, 25,830 x g) to pellet proteins. The supernatant was subsequently processed for metabolomics as described below. The protein pellets were frozen at −80 °C until ready for processing. The protein pellets were resuspended in 8 M urea/50 mM HEPES (pH 8.5). Tissues were homogenized in a TissueLyser II for 4 cycles at 29 Hz (Qiagen Hilden, Germany). Cells were briefly sonicated to solubilize the proteins. The lysate was centrifuged (16,000g, 15 min) to remove cellular debris. Proteins were reduced with dithiothreitol (5 mM, 56°C, 30min) and alkylated with iodoacetamide (15 mM RT, 30 min in the dark). Excess iodoacetamide was quenched with dithiothreitol (5 mM, room temperature, 30 min in the dark). The protein amount was quantified using Bradford (Pierce) and an aliquot of 50μg protein was digested using Lys-C (25 °C, 15 h) in a buffer comprising 50 mM HEPES (pH 8.5)/2 M urea. Following protein digestion, samples were acidified to a final concentration of 0.1% Trifluoroacetic acid and desalted using Oasis HLB 96-well uElution plate (Waters). Peptides were eluted with 50% acetonitrile/0.1% formic acid and dried overnight under vacuum at 30 °C (Labconco CentriVap Benchtop Vacuum Concentrator, Kansas City, Mo). Dried peptides were resuspended in 0.1% trifluoroacetic acid and fractionated using HPRP (High pH Reversed-Phase) according to the manufacturer’s instructions (Pierce). Four peptide fractions were collected per sample which were eluted with 7.5, 12.5, 17.5 and 50% acetonitrile/0.1% triethylamine. Samples were dried down under vacuum and reconstituted in 4% acetonitrile/5% formic acid for LC-MS/MS analysis.

Peptides were analyzed on an Orbitrap Fusion Lumos mass spectrometer (Thermo Fisher Scientific) coupled to an Easy-nLC (Thermo Fisher Scientific). Peptides were separated on an Aurora UHPLC emitter column (75 μm internal diameter, 25 cm long, pre-packed with C18 resin, 1.6 μm; IonOptiks). The total LC-MS run length for each sample was 85 min comprising a 70 min gradient from 6 to 25% acetonitrile in 0.125% formic acid. The flow rate was 300 nL/min and the column was heated to 60 °C.

Data-dependent acquisition (DDA) mode was used for mass spectrometry data collection. A high resolution MS1 scan in the Orbitrap (m/z range 375-1,540, 120k resolution, AGC 4 × 10^5, max injection time 50 ms, RF for S-lens 30) was collected. Dynamic exclusion was modified to a 60 second duration and excluded ion after 1 time. The MS2 scan was performed in the quadrupole ion trap (HCD, AGC 1 × 10^4, HCD collision energy 30%, max injection time 35 ms). For the LC-MS/MS runs on the cell lysates, the data were acquired in Profile mode.

### Quality Control for Peak Interference

A downside to labeling many amino acids is that the signal intensity tends to be distributed across a larger number of isotopologues, which increases the opportunity for a single interfering ion peak to distort the entire set of proportions. For this reason, quality control filters are commonly used in algorithms for analyzing D_2_O turnover data, even though they often result in a substantial loss of useable data. Our data confirm the need for an algorithm to remove interference, but we have also seen the dangers associated with biased filtering approaches.

Since our framework creates theoretical boundaries for each observation (the proportions must fall between the initial and final states) it is tempting to use these boundaries for quality control filtering. However, we have found that this can result in a severe underestimation of the experimental variability. As can be seen by a few simple simulations, valid observations frequently fall outside theoretical boundaries (Figure 2b). Worse still, by forcing or filtering for only observations within theoretical boundaries, we would be biasing our results towards the center of the sampling time window. Yet, without using the boundaries at all, the identification of outliers becomes very difficult. With these considerations in mind we deploy three filters.

The first filter only considers the labeled peptide prevalence. When the prevalence is less than a user-specified cutoff (10% by default) we remove the observations. As emphasized throughout the manuscript and highlighted in Figures 2c and 2d, low prevalence peptides provide very little value and may complicate model fitting since the parameters are non-identifiable when prevalence equals zero.

The second filter does not make use of any peptide sequence information. After dividing through by the sum of X isotopomer intensities, we then find the binomial distribution with size X that minimizes the Euclidean distance from our observation. We expect that each observed distribution should be similar to a binomial since the underlying problem of adding isotopes at the elemental level describes a binomial process, with zero or one denoting the presence of a heavy element. If the absolute value of the difference between any of the observed isotopologue proportions and associated binomial probability exceed a user entered parameter, residCut, then the observation is discarded. By default we set residCut to 0.2.

The residual cutoff from the binomial fit should remove most of the errant peaks that distort the shape of the distribution. However, it is still possible for the shape to be approximately correct while the observations remain impossibly far away from the theoretical boundaries. To avoid this case we create a second cutoff for the minimum absolute difference between either boundary and each binomial probability. If this difference exceeds the boundCut parameter (0.1 by default) then we remove the observation.

Through trial and error and visual inspection, we believe that these parameters are largely successful at removing interference without artificially reducing our estimate of experimental error. All analyses of real data described in this manuscript used the default filters just described.

### SILAC Validation (Experimental Methods)

For the SILAC validation experiment, growth media were prepared from a powdered DMEM high glucose base lacking lysine (AthenaES). For SILAC conditions, 13C6 15N2 L-lysine-2HCl (Thermo Fisher) was supplemented to 181 mg/L. For non-SILAC conditions, L-lysine-2HCl (Thermo Fisher) was supplemented to 175 mg/L. For D_2_O conditions, 99.9% D_2_O (Sigma) was used in the reconstitution of the powdered base, to a final D_2_O concentration of 8%. All media were supplemented with dialyzed FBS (Thermo Fisher) to 10% v/v.

HEK293T cells were grown in unlabeled media in 10 cm plates for all conditions of the experiment. At t = 0, existing growth media were replaced with media containing either heavy lysine or 8% D_2_O. Media in plates were further refreshed at t = 1 and t = 4 hours. At t = 0, 0.5, 1, 2, 4, 8, 24 and 48 hours, 3 replicate plates were harvested from each of the heavy lysine and 8% D_2_O conditions. To maintain sample integrity, all harvesting steps were performed on an aluminum block cooled to −20°C. Plates were washed with cold PBS. 3.5 mL of 80% methanol at −20°C was added to each plate and cells were scraped into a 5 mL Eppendorf Lo-Bind tube. Samples were stored at −20°C in preparation for metabolomic and proteomic workflows.

### In vivo protein turnover in naked mole-rats and C57BL/6 mice

Animal use and experiments were approved by the Buck Institute institutional animal care and use committee (IACUC) protocol number A10208. In total, 18 young C57BL/6 mice (11-13 weeks old, virgin males) and 18 young naked mole-rats (25-27 months old, non-breeding males) were used for this study. The selected age groups comprised young, healthy adults from both species that were physiologically age-matched (approximately 5-6% of observed maximum life span). Mice were purchased from the Jackson Laboratories (Bar Harbor, ME) and maintained in the vivarium for at least two weeks prior to use; naked mole-rats were from the captive colonies of Rochelle Buffenstein^36^.

For protein turnover studies, animals were injected intraperitoneally with a bolus of 99.9% D_2_O (1 ml/100 g body weight; Sigma Aldrich #151882) twice on the day of initiation of the experiment (Day 0; injections at 0 h and again at 8 h). Following this, the animals received a daily maintenance dose of D_2_O (0.5 ml/100 g body weight) by intraperitoneal injection. No dietary changes were made i.e. mice were allowed *ad libitum* access to (unlabeled) drinking water and chow, and naked mole-rats had *ab libitum* access to food (which is also their source of water). On days 1, 2, 4, 8 and 16, three animals from each species were anesthetized with isofluorane and euthanized by cardiac puncture. Following this, liver tissue was promptly harvested, cut into small pieces on ice and snap-frozen in liquid nitrogen. The snap-frozen tissues were subsequently used for metabolomics and proteomics experiments as described above. Note that sacrificed animals did not receive the maintenance dose of D_2_O on the day of sacrifice. Euthanization and organ collection was performed between 8 am and 11 am.

### Comparison of CIDS/BDEMS to existing proteomics tools

We sought to evaluate the performance of our BDEM-CIDS approach to estimate protein turnover. We generated a dataset of HEK293 human HeLa cells, and subjected these cells to isotopically labeled water containing 8% D2O. 3 extractions were collected after each of 4, 8, 24, and 48 hours following introduction of D2O. To compare the efficacy of our approach, we conducted a benchmark SILAC experiment and calculated protein turnover from these SILAC results as described below. We also evaluated two alternative existing tools for estimating protein turnover in D2O experiments, DeuteRater^19^ and d2ome^20^.

We retrieved the DeuteRater source code from https://github.com/JC-Price/DeuteRater. To execute this software, we followed the instructions provided in the user’s guide. In doing so, we uncovered a number of incompatibilities in the code that prevented successful execution. We corrected these issues and offer our modified source code in the forked repository https://github.com/PMSeitzer/DeuteRater. DeuteRater requires as input a list of identified peptides and proteins. We reformatted search results obtained by running both Comet and our CIDS approach into a DeuteRater-readable format (supplemental scripts X and Y). DeuteRater returned protein turnover rate estimates for all identified peptides and proteins associated with both Comet and CIDS, which we collected and analyzed further in a custom analysis script (script Z). Deuterator half-lives were taken from the output file named Final_Combined_Rates.csv. These results tables have been included in the supplementary files.

We downloaded the d2ome executables from https://github.com/rgsadygov/d2ome. d2ome is designed to intake raw, unsearched mzML files into a Mascot server, and calculate protein turnover rates based these search results. In our case, we were interested in evaluating our search engine results instead of Mascot, and so we created custom scripts to reformat our search results into mzIdentML files resembling the output of Mascot searches (Scripts X1, Y1). We produced two sets of results, one corresponding to Comet search results, the other to CIDS, and applied d2ome to both sets. Rates were taken from the output file named “Analyzed_Proteins_and_Their_Rates.csv” and were converted to half-lives.

In total, we generated protein turnover estimates for our data using eight different approaches: for each of SILAC, DeuteRater, d2ome, and BDEM, a regular Comet search and our CIDS search algorithm. To compare these approaches, we first reduced the datasets to overlapping proteins quantified in all methods. For each remaining protein we then subtracted the SILAC half-life estimate from each of the other results. These differences are the errors plotted in Figure 4c.

## Data availability

RAW files and other data tables will be made available at the time of publication.

## Code availability

Code for generating CIDS spectra and modeling protein turnover can be found at https://github.com/calico/D2O

## Contributions

JJO, YW, CS, FM and BB designed the in vitro validation experiment. The experiment was carried out by YW, NH and BB. The murine experiments were designed and implemented by BMM, GK, BMM, MK, CMS and NH. The NMR experiment was designed by JJO, VN, MS, FM, BB and RB. The experiment was performed by VN, NH, CMS, MS and RB. New algorithms were designed by JJO and VJ. JJO, PS, RR and AG developed software. JJO, VN, FM, AB, BB and RB wrote the bulk of the manuscript, while all authors contributed to and approved the final version.

